# Human cerebrospinal fluid induces neuronal excitability changes in resected human neocortical and hippocampal brain slices

**DOI:** 10.1101/730036

**Authors:** Jenny Wickham, Andrea Corna, Niklas Schwarz, Betül Uysal, Nikolas Layer, Thomas V. Wuttke, Henner Koch, Günther Zeck

## Abstract

Human cerebrospinal fluid (hCSF) have proven advantageous over conventional medium when culturing both rodent and human brain tissue. Increased excitability and synchronicity, similar to the active state exclusively recorded *in vivo*, reported in rodent slice and cell-cultures with hCSF as recording medium, indicates properties of the hCSF not matched by the artificial cerebrospinal fluid (aCSF) commonly used for electrophysiological recording. To evaluate the possible importance of using hCSF as electrophysiological recording medium of human brain tissue, we compared the general excitability in *ex vivo* human brain tissue slice cultures during perfusion with hCSF and aCSF. For measuring the general activity from a majority of neurons within neocortical and hippocampal human *ex vivo* slices we used a microelectrode array (MEA) recording technique with 252 electrodes covering an area of 3.2 x 3.2 mm^2^ and a second CMOS-based MEA with 4225 electrodes on a 2 x 2 mm^2^ area for detailed mapping of action potential waveforms. We found that hCSF increase the number of active neurons and the firing rate of the neurons in the slices as well as increasing the numbers of bursts while leaving the duration of the bursts unchanged. Interestingly, not only an increase in the overall activity in the slices was observed, but a reconfiguration of the network functionality could be detected with specific activation and inactivation of subpopulations of neuronal ensembles. In conclusion, hCSF is an important component to consider for future human tissue studies, especially for experiments designed to mimic the *in vivo* situation.

## Introduction

The brain, with all its neurons, glia cells and blood vessels, is cushioned and protected by a colourless fluid called cerebrospinal fluid (CSF). The CSF can be found in the space between the brain surface and the scull, the subarachnoid space, in all the ventricles as well as in and around the spinal cord. It is a fluid characterised by a low percentage of protein and high percentage of salts that is in constant contact with the interstitial fluid of the parenchyma, the fluid located between neurons and glia cells all over the brain. The close connection with the interstitial fluid gives the CSF a potential to act as a vessel for different neuromodulatory signals (Agnati et al., 1986; Agnati, Guidolin, Guescini, Genedani, & Fuxe, 2010) and experimental findings summarised in the review by Bjorefeld et al 2018 (Bjorefeldt, Illes, Zetterberg, & Hanse, 2018) show that the human CSF (hCSF) can influence the function of neurons in the both hippocampal slices and cortical neuronal cultures from rat. In general, all electrophysiological experiments performed using either rodent brain tissue slices or human brain slices from resected tissue use a solution called artificial CSF (aCSF) for perfusion of the tissue during the experiment. The aCSF is carefully made with several different salts together with glucose to mimic the components in CSF. But the CSF also contains a number of neuromodulators such as neurotransmitters, neuropeptides, neurosteroids, purines and endocannabinoids which is not present in the aCSF (Skipor and Thiery, 2008) (Bjorefeldt et al., 2018). When the hCSF was compared to the aCSF in experiments with rat hippocampal slices the spontaneous firing of action potentials increased in both hippocampal and cortical slices (Bjorefeldt et al., 2015). The activity mimicked what is usually only seen during recordings *in vivo* and activation of G-protein coupled receptors seemed to be the reason for the enhanced activity. Clear advantages of using hCSF as culturing medium for human organotypic brain slices has previously been shown by us. Experiments with cortical human brain slices, acquired from resected human brain tissue removed as part of treatment, resulted in healthier slices over both short and long incubation times when using hCSF compared to conventional culturing medium (Schwarz et al., 2017). Using resected human brain tissue as an *ex vivo* platform for modelling diseases or to study healthy brain activity provides huge translational advantages and several insights in healthy and disease related mechanisms have already been gained from this tissue (Dossi et al., 2018) (Mansvelder et al., 2019). The major limitation of the human tissue *ex vivo* platform is the lacking of directly comparable *in vivo* result. It is therefore of great importance to design the *ex vivo* experiments to mimic the *in vivo* situation as much as possible.

It is now known that human tissue survives better when cultured with hCSF compared to conventional cell culture medium but if the hCSF affect the function of the human neurons and level of activity is still an open question. The aim of our present study is to find out whether perfusion of hCSF affect the neuronal activity, on a single cell and network level, compared to perfusion with the commonly used aCSF. We hypothesised that, just as was seen in the rat study, the general activity in the neurons will increase with perfusion of hCSF compared to aCSF. We perform extracellular recordings using high-density micro-electrode arrays (MEA) to allow for analysis of the response from both individual neurons, and how their activity contribute to the overall network activity (Ferrea et al., 2012; Gong et al., 2016; Zeck, Jetter, Channappa, Bertotti, & Thewes, 2017).

## Materials and methods

Human hippocampal and cortical organotypic slice cultures were prepared from resected tissue obtained from patients undergoing epilepsy surgery. For this study, we collected and included data of 5 patients. All patients were surgically treated for intractable epilepsy. Approval (# 338/2016A) of the ethics committee of the University of Tübingen as well as written informed consent was obtained from all patients, allowing spare tissue from resective surgery to be included in our study. Hippocampus was carefully micro dissected and resected *en block* to ensure tissue integrity, directly transferred into ice-cold aCSF (in mM: 110 choline chloride, 26 NaHCO3, 10 D-glucose, 11.6 Na-ascorbate, 7 MgCl2, 3.1 Na-pyruvate, 2.5 KCl, 1.25 NaH2PO4, und 0.5 CaCl2) equilibrated with carbogen (95% O2, 5% CO2) and immediately transported to the laboratory. Tissue was kept submerged in cool and carbogenated aCSF at all times. The tissue was trimmed to give a flat glue-surface in the coronal plane, glued onto the slicing platform (Figure 1A) and then sliced in to 250 µm thick slices using a Microm HM 650V vibratome (Thermo Fisher Scientific Inc). Subsequently, the slices were transferred onto culture membranes (uncoated 30 mm Millicell-CM tissue culture inserts with 0.4 µm pores, Millipore) and kept in six well culture dishes (BD Biosciences) as shown in Figure 1A. The plates were stored in an incubator (ThermoScientific) at 37°C, 5% CO2 and 100% humidity. For electrophysiological recordings slice cultures were transferred to the MEA-recording chamber. We received and pooled hCSF of several patients with normal pressure hydrocephalus (NPH) who needed to undergo a tap test either by lumbar puncture or lumbar drain as part of the diagnostic workup. Approval of the ethics committee of the University of Tübingen as well as written informed consent was obtained from all patients. It is well established and known from daily clinical practice that hCSF of NPH patients exhibits physiological hCSF parameters (lactate, glucose, cell count, protein levels) see (Schwarz et al., 2017), which are undistinguishable from the ones of healthy individuals (Bjorefeldt et al., 2015). The hCSF was centrifuged at 4000 rpm at 4°C for 10 minutes and the supernatant was collected and stored at −80°C.

**Figure 1:**
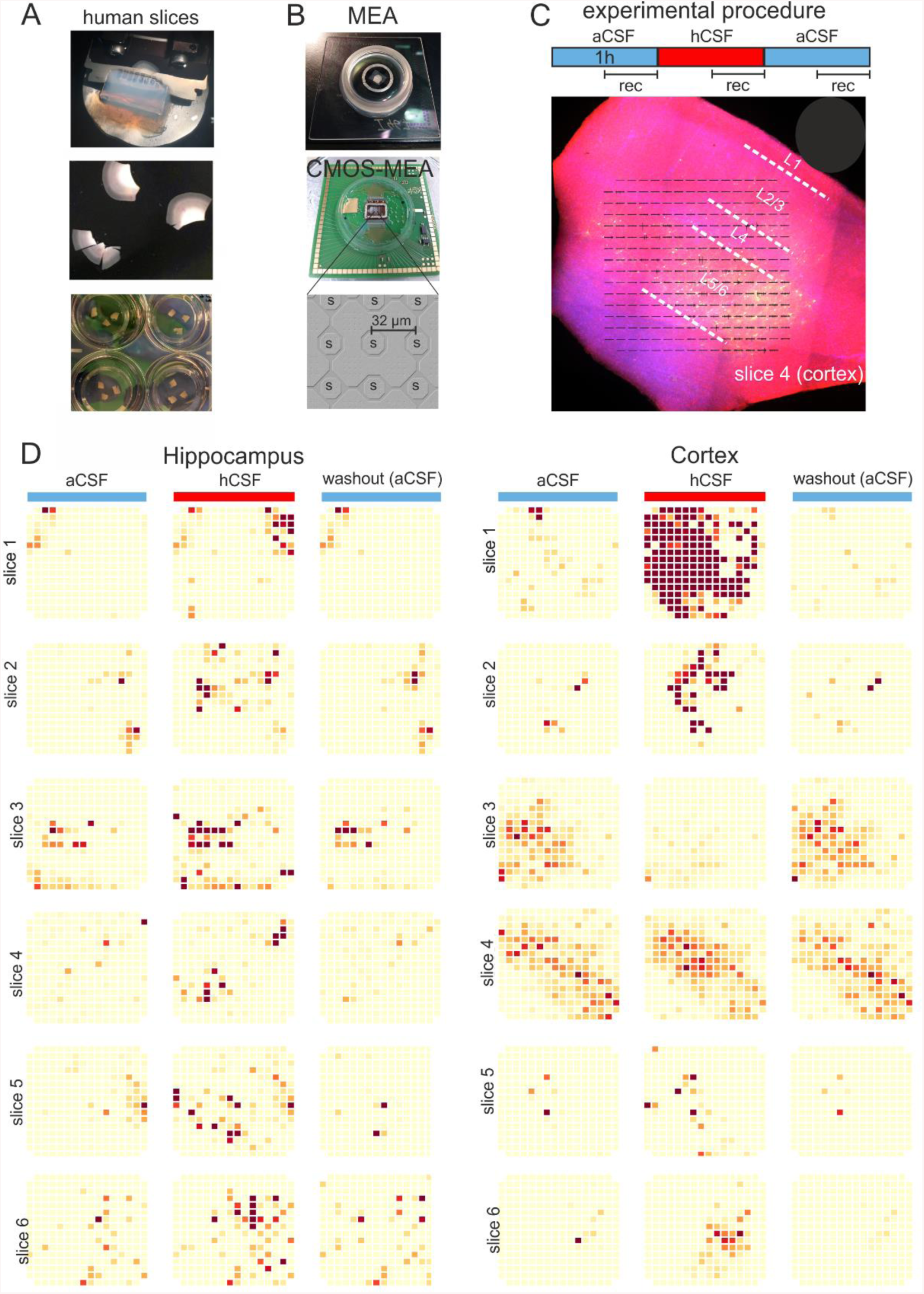
Experimental design and overview of activity: (A) The brain tissue is surgically resected *en block* and the tissue block is cut into 250 µm slices using a vibratome. The slices are incubated on membrane-insets in 6-well plates with hCSF as medium for up to two weeks. (B) Pictures of the 256-MEA chip (top), the CMOS-MEA chip (middle) and the sensor spacing of the CMOS-MEA. (C) Experimental protocol. Recording was performed continuously for 30 minutes in aCSF / hCSF / aCSF. (D) Qualitative overview over twelve slices recorded using MEA 256. When hCSF is washed in the general activity of the slice increased with new areas of the slice active and increased number of spikes presented here as heat-maps from three hippocampal and three cortical slices. Each electrode is represented by a small square and darker colour indicates higher spike rate. Activity is normalized for each slice to the maximum spike rate per electrode in aCSF.

### Micro-electrode array recordings

The majority of slices were transferred from the incubation chamber to a 256-electrode MEA (Multi Channel Systems MCS GmbH) with continuous perfusion of aCSF heated to 34°C (Figure 1B). The aCSF contained (in mM) 118 NaCl, 3 KCl, 1.5 CaCl_2_, 1 MgCl_2_, 25 NaHCO_3_, 30 D-glucose, and equilibrated with carbogen (95% NaH_2_PO_4_, and 30 O2–5% CO_2_, pH 7.4) in a recycling system. The electrode spacing was 200 µm with the total recordings area of ∼ 3.2 x 3.2 mm^2^. The recordings were made with a sampling rate of 25 kHz using the MC Rack software (Multi Channel Systems MCS GmbH) and each condition, aCSF, hCSF and rewash of aCSF was performed for 1 hour. To avoid analysing data from the transition-time of one medium to the next the analysis window was defined as the last half hour of each condition.

A subset of slices (n = 3) was investigated using a CMOS-based MEA setup (CMOS MEA 5000, Multi Channel Systems MCS GmbH). The recording chip comprised 4225 sensor sites (arranged in 65 x 65 sensor lattice) with an electrode pitch of 32 µm thus covering a recording area of ∼ 2 x 2 mm^2^ (Figure 1B) Recordings were performed at 20 kHz using CMOS MEA Control software (Multi Channel Systems MCS, version 2.2.0).

### Data analysis and statistics

The analysis of recordings from 256-MEA was performed using Python (version 3.6, including the libraries numpy, scipy). Recordings were bandpass - filtered (150 – 5000 Hz, butterworth 2nd order) and spikes were detected as described (Quian Quiroga et al., 2004) using a 1 ms post-spike dead-time. The firing rate for each electrode was calculated based on the number of threshold crossings within the recording time (30 minutes). To reduce the number of false positive detection of active electrodes only electrodes with more than 90 identified spikes in the thirty-minute long recording were considered as active. Thus the average firing rate per slice comprised only electrodes with a firing rate higher 0.05 Hz. The colour scale of the heatmaps displaying the average firing rate in each slice in Figure 1D were normalized to the maximum firing rate detected in the aCSF (first condition) for each slice.

### Single channel bursts

Single channel bursts were defined as having at least 3 consecutive spikes with a maximum of 100 ms inter-spike interval. This definition was adapted using previous reports (Chen et al., 2009) and the joint inter-spike-interval histogram of individual electrodes. Increasing or decreasing firing rate between aCSF and hCSF was inferred if the relative change exceeded 50 percent. Electrodes with less than 50 percent change were considered stable.

### Population bursts

The population bursts were defined as described previously (Koch et al., 2019). In brief, for the identification of population bursts, all detected spikes were merged in consecutive non-overlapping 5 ms bins. This raw population firing rate was smoothed using a normalized Gaussian kernel with standard deviation 100 ms. Network burst onset was determined when this smoothed population firing rate exceeded the slowly varying 1-second moving average. Very low-firing population bursts were discarded, if their peak firing rate did not exceed 10% of the average of the top five peaks. Population burst detection was eventually checked by visual inspection to avoid detection of occasional artefacts present on all electrodes.

### CMOS-MEA

Analysis of the recordings using the CMOS-MEA were performed using CMOS MEA Tools, a software implementing the cICA algorithm (Leibig et al., 2016). Only well-isolated single units were considered (criterium: IsoBG > 5). Due to computational constrains, continuous recordings with the CMOS-MEA were, in the present study, limited to 2 minute-long files.

All statistical testing was done with Prism7 software (Graphpad) and differences between groups were considered significant when p < 0.05.

## Results

Seven hippocampal slices from two hippocampectomy procedures and eight cortical slices from three temporal lobe resections resections were collected for extracellular recording using 256 – MEAs (six hippocampal slices, six cortical slices) and CMOS-based MEA (one hippocampal slice and two cortical slices). After 7 - 14 days in vitro culture using hCSF the slices were placed in the MEA recording chamber, perfused with aCSF for an acclimation period of 30 minutes. After acclimation, aCSF was perfused for an additional hour (we name this period *aCSF)* followed by one hour of hCSF perfusion (we name this period *hCSF*) and then a final hour with aCSF (we name this period *washout*), see Figure 1C.

### Human CSF induces a general increase in neuronal activity

To quantify the excitability level of the 12 slices on the 256 - MEAs we first extracted the number of active electrodes (as defined in the Method section) for each condition, showing 60.67 ± 18.15 number of electrodes active during *aCSF*, 95.42 ± 15.58 during *hCSF* and 59.08 ± 20.69 during *washout.* The data from the three conditions was tested with the paired, non-parametric Friedman test and a significant difference was found (p = 0.0062) and Dunn’s multiple comparison test found the increase in number of active electrodes when comparing *aCSF* and *hCSF* significant (p = 0.0239) as well as the decrease in number of active electrodes when comparing *hCSF* and *washout* (p = 0.0128) while the number of active electrodes remained stable between *aCSF* and *washout* (p > 0.9999), altogether demonstrating reversibility of the *hCSF* mediated activity increase (Figure 2A). We next analyzed the overall spike rate in each slice and found the average spike rate to be 0.95 ± 0.50 spikes/s during *aCSF*, 1.76 ± 0.43 spikes/s during *hCSF* and 1.08 ± 0.57 spikes/s during *washout*. The average spike-rate in each slice increased when washing in hCSF and then decreased when switching back to the aCSF again during the *washout*. The data from the three conditions was tested with the paired, non-parametric Friedman test, a significant difference was found (p = 0.0014) and Dunn’s multiple comparison test found the increase in firing rate seen when comparing *aCSF* and *hCSF* significant (p = 0.0016) as well as the decrease between the *hCSF* and the *washout* (p = 0.0239) while the firing rate remained stable between *aCSF* and *washout* (p > 0.9999) (Figure 2A).

**Figure 2:**
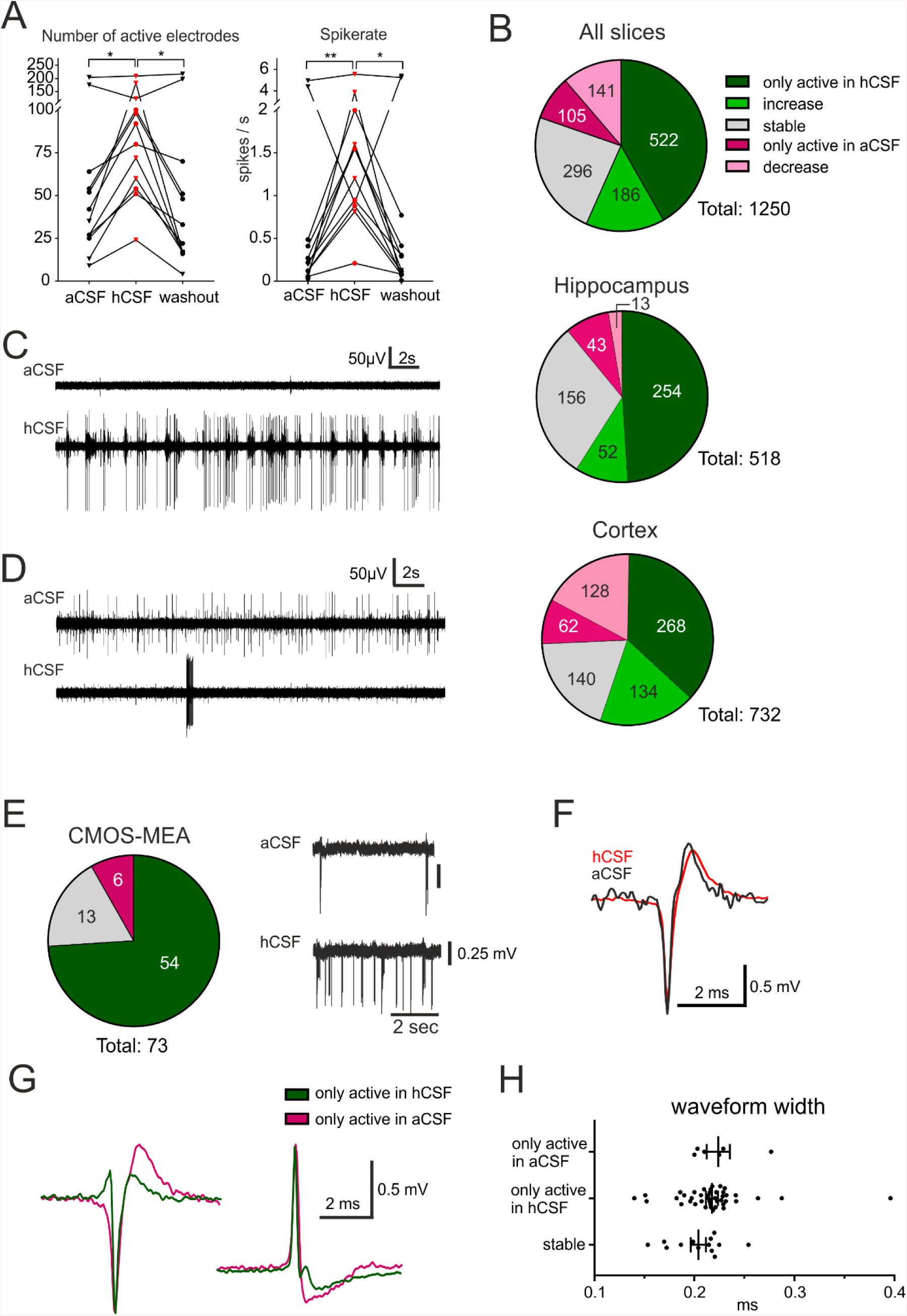
hCSF induces a general increase in neuronal activity and specific activation and inactivation of subpopulations of neurons. (**A)** The general activity measured by number of active electrodes and spike-rate, with an average of each hippocampus slice represented by a circle and each cortex slice as a tringle, is increasing when hCSF is washed in and then decreasing again when hCSF is washed out. The average number of electrodes and spike-rate in one cortical slice did not increase but decreased instead (see Figure 1D, cortex slice 3). **(B)** We analysed the activity in individual 256-MEA recording electrodes from all slices and grouped the electrodes based on activity: only active in *hCSF*, increased spike-rate in hCSF, decreased spike-rate in *hCSF*, only active in *aCSF* and stable (including the electrodes that have an increasing or decreasing spike-rate less than 50%). The majority of the electrodes only recorded spikes in hCSF or measured an increased spike-rate in hCSF, 522 and 186 electrodes respectively of a total of 1250 electrodes. Interestingly, 105 electrodes only recorded activity in *aCSF* and 141 electrodes recorded a decreased spike-rate in *hCSF* compared to *aCSF*. Examples of electrode recordings revealing increased activity (**C)** and decreased activity (D) when *hCSF* is washed in. The three conditions (*aCSF, hCSF* and *washout*) were tested with the paired, non-parametric Friedman test followed by Dunn’s multiple comparison test, *p < 0.05; **p < 0.01. Changed excitability was also detected in extracellular recordings using the CMOS MEA. (**E**) Overview of identified single-unit activity (n = 73 cells in 3 slices) with extracellular voltage recorded by a selected sensor showing the increased excitability of one cell in *hCSF* compared to *aCSF*. Selected extracellular waveforms of cells active in **(F)** both conditions, **(G)** in *hCSF* and in *aCSF* only. **(H)** The waveform width was not different between the spikes recorded from neurons only active in *aCSF*, only active in *hCSF* or active in both (stable).

Next we performed a subgroup analysis and assessed the number of active electrodes and the firing rates separately for hippocampal and for cortical recordings. While in the hippocampus subgroup we found the same significant differences as described for pooled data from both groups, no significant differences were found in the cortex subgroup (Table 1 for mean ± SEM and p-values). The data from two of the slices is clearly going against the trend, see Figure 2A. To better understand why some of the slices displayed such a clear deviation from the rest of the group we performed a deeper analysis of the activity in each electrode.

### Specific activation and inactivation in subpopulations of neurons

Further analysis of the firing rate recorded in each individual electrode detected three distinct responses to washing in of *hCSF*; i) decreased firing rate, ii) increased firing rate and iii) no change in firing rate. An average of 9.3 ± 1.9 electrodes recorded a decreased firing rate in each slice and 5.2 ± 1.4 of them per slice recorded no spikes during *hCSF*. The number of electrodes measuring an increase in spike rate was on average 58.7 ± 8.9 per slice and 40.3 ± 4.8 of them recorded no spikes during *aCSF*. On average, 11.3 ± 2.6 electrodes recorded a change (increase or decrease) in firing rate that was less than 50% and was considered stable. These data indicate that individual neurons or subnetworks in the slices respond differently when *hCSF* is washed in, with some neurons decreasing action potential firing but with the majority of neurons responding with increased action potential firing (Figure 2B-D).

For three additional slices we performed a detailed analysis of the specific activation based on identified cellular activity using CMOS-MEAs. Recordings were performed under similar experimental conditions as with the 256 –MEAs (Figure 1C, aCSF / hCSF / aCSF). In these recordings we identified 73 individual cells, while many electrodes recording multi-unit activity were removed from further analysis. The majority of identified cells (74%) was only active in *hCSF*, a small number (8%) was only active in *aCSF* and the remaining cells (18%) were identified as active in both *hCSF* and *aCSF* (Figure 2E). These results support our 256 – MEA findings that different subpopulations of neurons may respond distinctly to *hCSF* and *aCSF*.

Spike sorting provided the average extracellular waveform of each cell on multiple neighbouring electrodes. Searching for potential waveform differences between aCSF and hCSF we selected for each cell the electrode with the maximal amplitude. For cells identified as active in both aCSF and hCSF (n = 13) no change in the waveform could be observed between *aCSF* and *hCSF* (Figure 2F). However, when comparing the waveforms between cells the shapes were different, with most of them displaying a negative peak but some also positive peaks (Gold et al., 2009). The average width at half waveform peak was 0.21 ± 0.03 ms with all identified cells included and did not differ between cells only active in *aCSF* (0.22 ± 0.03 ms), cells only active in *hCSF* (0.21 ± 0.04 ms) or cells active in both *aCSF* and *hCSF* (0.20 ± 0.03 ms) (Figure 2G-H). We identified five cells active only in *hCSF*, which displayed a narrow waveform (< 0.16 ms), a potential indication for inhibitory neurons (Viskontas et al., 2007). In summary, the extracellular waveforms did not provide indications for differential effects of *hCSF* on neurons within human brain slice cultures.

### Human CSF induces an increase in burst-activity and a synchronisation shift

To further understand how neurons change their activity when hCSF is washed in we looked specifically at burst activity (defined in the Method section) on individual active MEA electrodes and found that the average number of bursts in each slice during *aCSF* was 11637 ±7616, during *hCSF* 16597 ± 9561 and during the *washout* 12258 ± 7839. The data from the three conditions were tested with the paired, non-parametric Friedman test, and a significant difference was found (p = 0.0052). Dunn’s multiple comparison test found the increase in number of bursts seen when comparing *aCSF* and *hCSF* significant (p = 0.0322) as well as the decrease between the *hCSF* and the *washout* (p = 0.0092) while the number of bursts remained stable between *aCSF* and *washout* (p > 0.9999). The duration of the detected bursts was, on average during *aCSF* 0.43 ± 0.06 s, during *hCSF* 0.64 ± 0.20 s and during *washout* 0.38 ± 0.07 s. The data for the three conditions was tested with the paired, non-parametric Friedman test and no significant difference was found (p = 0.5580). To evaluate the effect hCSF has on network activity, we continued by detecting the number of population bursts (defined in the Method section) and the number of cells participating in the detected population bursts. Population bursts were detected in nine (four hippocampal and five cortical slices) of the twelve slices with number of population bursts during *aCSF* 45 ± 29.8, during *hCSF* 190 ± 76.3 and during *washout* 33 ± 23.1. The three conditions were tested with the paired, non-parametric Friedman test, a significant difference was found (p = 0.0148) and Dunn’s multiple comparison test found the decrease in number of population bursts seen when comparing *hCSF* and *washout* significant (p = 0.0201) but *aCSF* compared to *hCSF* (p = 0.3765) as well as *aCSF* and *washout* (p = 0.7158) was found non-significant. The number of electrodes active in these population bursts was counted during *aCSF* to be 22 ± 6, during *hCSF* 84 ± 18 and during *washout* 32 ± 1. When testing the three conditions with the paired, non-parametric Friedman test a significant difference was found (p < 0.0001) and Dunn’s multiple comparison test found the increase in number of active electrodes during population bursts seen when comparing *aCSF* and *hCSF* significant (p = 0.0286) as well as the decrease between the *hCSF* and the *washout* (p = 0.0005) but the *aCSF* compared to the *washout* (p = 0.7158) was found non-significant. Looking at individual slices (individual points in the graph), see figure 3A, it becomes evident that the hippocampal slices (circles) did not show the same clear increase in number of population bursts during hCSF as the cortex slices (triangle). In the hippocampal slices we observed that in general the electrodes that recorded new activity in hCSF, record continuous spiking with no or little participation in population bursts, see figure 3B (hippocampal slice 5 heat maps in Figure 1). With the exception of one hippocampal slice where a completely new area (see Figure 1D, hippocampus slice 1, top right corner) becomes active during hCSF and is not engaged in the long population bursts but is having high frequency population burst separately from the rest of the slice, see figure 3C. In the cortex slices there is a clear shift towards higher level of synchronisation in the network. In several cortical slices the number of electrodes recording activity is not increasing much but the activity recorded is becoming highly synchronised with short population bursts engaging most of the network, see figure 3D for an example of this synchronisation shift observed in cortex slice 4 (heatmap in figure 1D). Even in the one slice where the number of active electrodes is decreasing in *hCSF*, cortex slice 3 (heatmap in figure 1D), the population bursts are appearing and most of the activity in the slice is synchronised during *hCSF*, see figure 3E.

**Figure 3:**
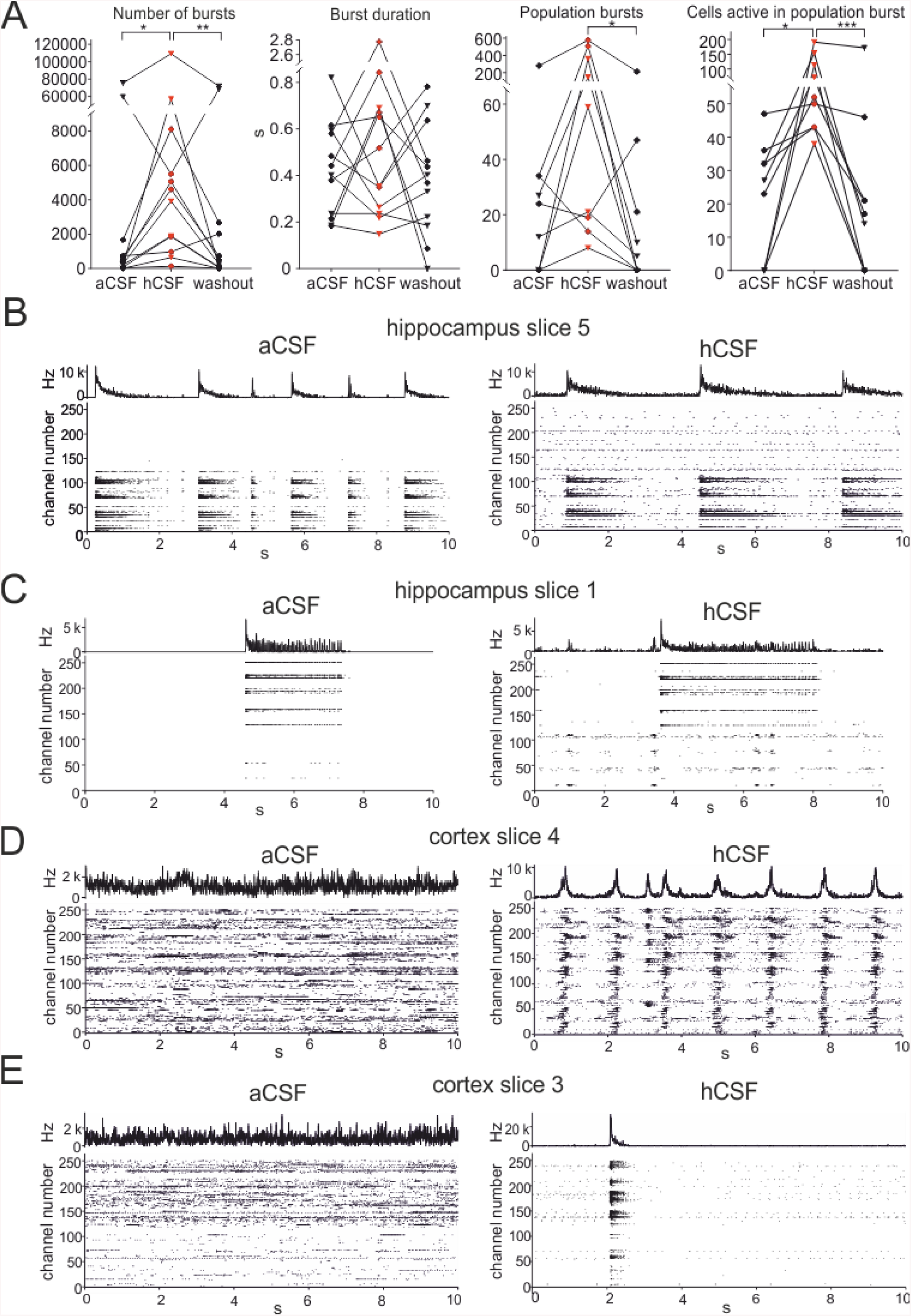
hCSF promotes synchronised network activity. (**A)** Bursts detected in single electrodes increase when hCSF is washed in and decrease back towards aCSF baseline levels when hCSF is washed out again (each hippocampus slice represented by a circle and each cortical slice as a tringle) (**A**) Burst durations do not change when hCSF is washed in. (**A**) Number of population bursts increased in all cortical slices (triangles) and in two of the hippocampal slices (circles) following hCSF wash in and then decreased in all slices but one (hippocampal) during the washout. The number of neurons (measured by number of active electrodes) participating in a population bursts is increased in all slices during hCSF and then decreased back toward baseline levels during washout. A 10 second representative raster plot with all electrodes included and summarised spike-rate trace above from: (**B**) hippocampus slice 5 (see heatmap in Figure 1D) showing that the activity of neurons becoming active in *hCSF* is in general not participating in the population burst but rather depicts continuous spike activity(**C**) hippocampus slice 1 (see heatmap in Figure 1D) showing a new area of the slices becoming active in *hCSF* with synchronised population bursts, (**D**) cortex slice 4 showing how non-synchronised activity recorded in *aCSF* becomes highly synchronised in *hCSF* across the slice and (**E**) cortex slice 3 (see heatmap in Figure 1D) shows that though the general number of spikes is reduced in *hCSF* a synchronisation shift is present and most spikes recorded are synchronised over the entire slice.

## Discussion

The activity of *in vitro* neuronal networks is highly dependent on the environment it is exposed to and is commonly experimentally measured in standard media or aCSF. In several recent studies the activity in rodent cells and networks was compared between aCSF and hCSF and in general an increase in the activity was detected (Bjorefeldt et al., 2015, 2018, 2019; Koch et al., 2019; Perez-Alcazar et al., 2016). In line with these studies we found that the general level of excitability increased in resected human hippocampal and neocortical slices when the slices were perfused with hCSF compared to the conventionally used aCSF, indicating that this effect is also preserved in human neurons and networks. The response when exposed to hCSF was rapid (present within minutes) and robust (lasting for the experimental protocol of one hour).

### hCSF and excitability

Using a 256-MEA we observed an increase in excitability by analysing number of active electrodes, spike-rate, number of bursts, burst duration, population bursts and number of electrodes active in population bursts. In a first step, we averaged these parameters for each slice recorded and in all slices but two the wash in of hCSF induced an increase in number of active channels, spike-rate and number of busts followed by a fast decrease back to baseline when hCSF was washed out and replaced by aCSF. As a previous study demonstrated in mouse and rat tissue, the general increase in excitability could not be explained by a simple increased level of potassium in the hCSF compared to the aCSF. The cortical slices had a clear shift from random spiking to very synchronised burst activity detected by numerous populations bursts and exemplified in figure 3D. In hippocampal slices the shift in synchronisation was not as clear as in the cortex slices but still changed towards an increased activity and in some slices towards a clear increase in synchronised activity, see figure 3B. In a recent study hCSF induced an increase in spontaneous gamma oscillations in CA1 and CA3 area of mouse hippocampal slices and the increase was shown to relay on muscarinic acetylcholine receptors (Bjorefeldt et al., 2019). The increase in bursts and population bursts induced by hCSF observed in our human tissue slices may indicate that the neurons usually engaged in oscillatory activity such as gamma oscillations more readily do so spontaneously when hCSF is present.

### Specific activation and inactivation of subpopulations of neuronal ensembles

Interestingly when analysing the activity recorded on each electrode more closely we found that the spike-rate in sub groups of neurons increased, did not change or decreased under the influence of hCSF. This result, together with the result that 42% of the electrodes only recorded activity in the presence of hCSF and 8% only recorded activity in the presence of aCSF, indicated not only an increase in the overall activity in the slices was observed, but a reconfiguration of the network with specific activation and inactivation of subpopulations of neuronal ensembles. One hypothesis explaining the specific activation and inactivation of different neuronal ensembles could be that an increased excitability among fast spiking interneurons, previously reported in rodent studies (Bjorefeldt et al., 2019), will in turn inhibit other neurons populations that would turn silent in hCSF. By characterising the cell-specific waveform of extracellularly recorded spikes and combining this with the spike-rate of the same cell it may be possible to discriminate between interneurons and excitatory neurons (Csicsvari et al., 1999). Interneurons have been characterized by narrower waveforms, higher spike rate and distinct bursting behavior as compared to principal neurons (Viskontas et al., 2007). As hCSF did not induce a change in the waveform compared to the waveform recorded in aCSF, measured from cells with spikes recorded in both aCSF and hCSF, the waveform width could be used to search for cell-type specificity. However, even if the 5 cells identified in hCSF with narrow width would be fast spiking interneurons the majority of cells (49) would have waveforms comparable to the average. Thus, we cannot conclude any cell-specific activation by *hCSF*.

Differences in calcium concentration have recently been shown between aCSF and hCSF followed by higher rates of spontaneous spikes recorded in rodent slices if the calcium concentration is not increased to the aCSF-level (Forsberg et al., 2018). This could be a possible explanation for the increase in excitability we have observed in the human tissue slices but future experiments specifically designed to test this will be needed before any conclusions can be made.

### Human tissue and hCSF

The resected human tissue is of great importance for translational research, especially for frequent neurological diseases such as epilepsy. The tissue provides a unique possibility to understand the physiological as well as pathological mechanisms in the human neuronal network circumventing the problem of poor translatability from animal models. Several studies using human tissue have already moved the research field forward providing new insights in pathophysiological mechanisms underlying temporal lobe epilepsy (Beck, Goussakov, Lie, Helmstaedter, & Elger, 2000) thereby not only helping to develop new antiepileptic drugs (Straub et al., 2001) but also increasing the general physiological understanding of cortical activity (Eyal, Mansvelder, de Kock, & Segev, 2014; M. B. Verhoog et al., 2013; Matthijs B. Verhoog et al., 2016)). Due to the limited access to resected tissue it is critical for results and studies to be compared between laboratories and different research groups working with this valuable tissue. Using standardised methods and solutions are key factors for enabling such comparisons, as recently pointed out (de la Prida and Huberfeld, 2019) the aCSF is as such a standardised solution. The results of the present study and others (Bjorefeldt et al., 2015; Bjorefeldt et al., 2019; Forsberg et al., 2018) show a clear difference in the neuronal firing behaviour when comparing aCSF and hCSF. To understand mechanisms during both physiological and pathological activity within the human brain network it is crucial to mimic the environment in the brain as closely as experimentally possible.

In conclusion, using hCSF instead of aCSF when recording from human tissue induces significant changes in the level of excitability. We demonstrate that the changes are complex and that the specific activation and inactivation of certain neurons cannot simply be explained by different effect on excitatory or inhibitory neurons. In the present study we have only revealed the major changes and highlighted the complexity, scratching the surface of the interesting effects hCSF has on individual neurons and neuronal networks in human tissue. Future studies are needed to fully understand which components of the hCSF induce the changes and how these affect individual neurons and neuronal networks we are studying.

## Acknowledgments

This work was supported by the German Ministry of Science and Education (BMBF, FKZ 031L0059A) and by the German Research Foundation (KO-4877/2-1 and KO 4877/3-1)

## Author contribution

JW and AC performed micro-electrode array recordings, NS, BU, NL, TW, HK and JW did tissue preparation and culturing, JW, AC, GZ did data analysis and writing. Proof reading and study design: all.

